# Plant growth promotion in *Oryza sativa* by *Bradyrhizobium sp.* isolated from the rhizosphere of *Aeschynomene indica*

**DOI:** 10.1101/2024.05.27.596129

**Authors:** Karl Joseph Samuel, R. Rajeswari

## Abstract

*Aeschynomene indica* is traditionally used as green manure in Tamil Nadu, India, to cultivate *Oryza sativa*. This plant contains *Bradyrhizobium sp*. in its root and stem nodules as PGPR. In this work, we isolated *Bradyrhizobium sp.* from the root nodules of *A. indica* using a selective medium and used it as a biofertilizer for *O. sativa*. The isolated microorganism was confirmed by various biochemical tests of Bergey’s Manual. Plant growth activity was measured in terms of plant height, plant diameter, leaf width, leaf length, grain weight, soil nitrogen fixation and nutritional value of the grains. Growth promotion of isolated *Bradyrhizobium sp*. was compared with commercial NPK fertilizers and commercial *Rhizobium* liquid fertilizer. Plants treated with *Bradyrhizobium sp.* provided a yield of 37.2 grams with average weight of 0.0182 g/grain while NPK treated plants had 38.4 g with average weight of 0.0175 g/grain. Total nitrogen fixed by *Bradyrhizobium sp.* (192 kb/ha) is almost similar to that of commercial NPK fertilizer (196 kg/ha). The nutritional value of the grains produced by *Bradyrhizobium* treatment (302 kcal) is higher than that of NPK fertilizer (286.12 kcal). Thus, free-living *Bradyrhizobium sp.* can be used as a biofertilizer for crops which does not have nodulation capability.

## Introduction

The blue planet is moving towards green with the help of sustainability. The use of artificial nitrogen-fixing agents such as the DAP is widely used in agriculture. The use of chemical fertilizers will improve the products in terms of quantity. The use of bio-based growth promoters is getting attention due to their cost and availability. This increases biological nitrogen-fixing agents like bacteria and other living organisms, converting nitrogen into ammonia (Zahran 1999). The bacteria which have nitrogenase enzyme involves in nitrogen fixation (Kim and Rees 1994). The biological nitrogen fixation process was done by mixing seeds and irrigation (Peoples, Herridge, and Ladha 1995).

In traditional cultivation of rice *(Oryza sativa)* in Tamil Nadu, India, the *Aeschynomene indica* plants belonging to the family *Fabaceae* are grown on the field before rice. During preparation of the field for planting rice seedlings, the previously grown *A. indica* plants are folded in to the soil and used as green manure for growing Rice. *A. indica* has stem nodules containing slow growing rhizobacteria *Bradyrhizobium sp* (Alazard 1985). It has heavy metal tolerance and free-living ability (Wani, Khan, and Zaidi 2007) with photosynthetic ability in some strains. They exhibit a slow growth in Yeast Extract Mannitol Agar (YEMA) with opaque, circular, and convex colonies (Jordan 1982). Photosynthetic *Bradyrhizobium* species have also been isolated from stem nodules of various *Aeschynomene* plants (van Berkum, Tully, and Keister 1995; Molouba et al. 1999). Significant plant growth promotion activity is observed in the plants inoculated with *Bradyrhizobium sp.* This activity is attributed to the gibberellin like substance (Dobert, Rood, and Blevins 1992) and auxin (Cassán et al. 2009). Successful studies have been conducted for using *Bradyrhizobium sp.* as PGPR for non-legumes has been conducted using Radish (*Raphanus sativus L.*) plants (Antoun et al. 1998).

Based on the research conducted by various authors, we believe that the free living *Bradyrhizobium sp.* can be used as a plant growth promotor for the non-legumes and plants without nodulating capability. This study aims to use free living *Bradyrhizobium sp.* isolated from the rhizosphere of *Aeschynomene indica* to promote nitrogen fixation and plant growth in *Oryza sativa*.

## Materials and Methods

### Isolation and Growth of PGPR from Aeschynomene indica

Seeds of Aeschynomene indica plants were bought from a local seed shop, and the plants were grown in pots. *A. indica* plants were grown for about two months. When the plants reached a height of 1.5 𝑚, they started flowering. First, the soil near the rhizosphere was collected. 5 𝑔 of soil was mixed with 25 𝑚𝑙 of distilled water, diluted by serial dilution, and then spread plated on *Bradyrhizobium* Selective Medium (BRSM).

#### Selective Medium

A selective medium for the growth of *Bradyrhizobium* was proposed by Gault and Schwinghamer (1993) and Tong and Sadowsky (1994). Selectivity of these media is determined by the antibiotic resistance and heavy metal tolerance of *Bradyrhizobium* strain, respectively.

The selective medium was prepared by modifying the BJSM proposed by Tong and Sadowsky (1994). The AG medium used by Tong and Sadowsky was a growth medium to support *Bradyrhizobium japonicum*. We replaced AG Medium with Yeast Extract Mannitol Medium, which is a standard growth medium for Rhizobacteria. Among all heavy metals tested by Tong and Sadowsky (1994), tolerance of Cobalt and Zinc showed a significant difference among the strains of *Rhizobium* and *Bradyrhizobium*. *Rhizobium* strains were tolerant to both 𝐶𝑜𝐶𝑙_2_ and 𝑍𝑛𝐶𝑙_2_ up to 40 𝜇𝑔/𝑚𝑙 while, *Bradyrhizobium* strains were up to 80 𝜇𝑔/𝑚𝑙. The concentration of Heavy metals in the selective medium is set to70 𝜇𝑔/𝑚𝑙 a concentration in which *Rhizobium* strains cannot grow while *Bradyrhizobium* can grow. Cycloheximide is used as a fungal inhibitor as suggested by Tong and Sadowsky. To inhibit the growth of Gram-positive organism, 0.5 𝜇𝑔/𝑚𝑙 of Brilliant Green (BG) was added to the medium.

BG and Cycloheximide were added to the sterile molten medium in an aseptic manner. The final composition of the AINS-03 Selective medium (BRSM) developed was as follows in table 1.

**Table 1.**
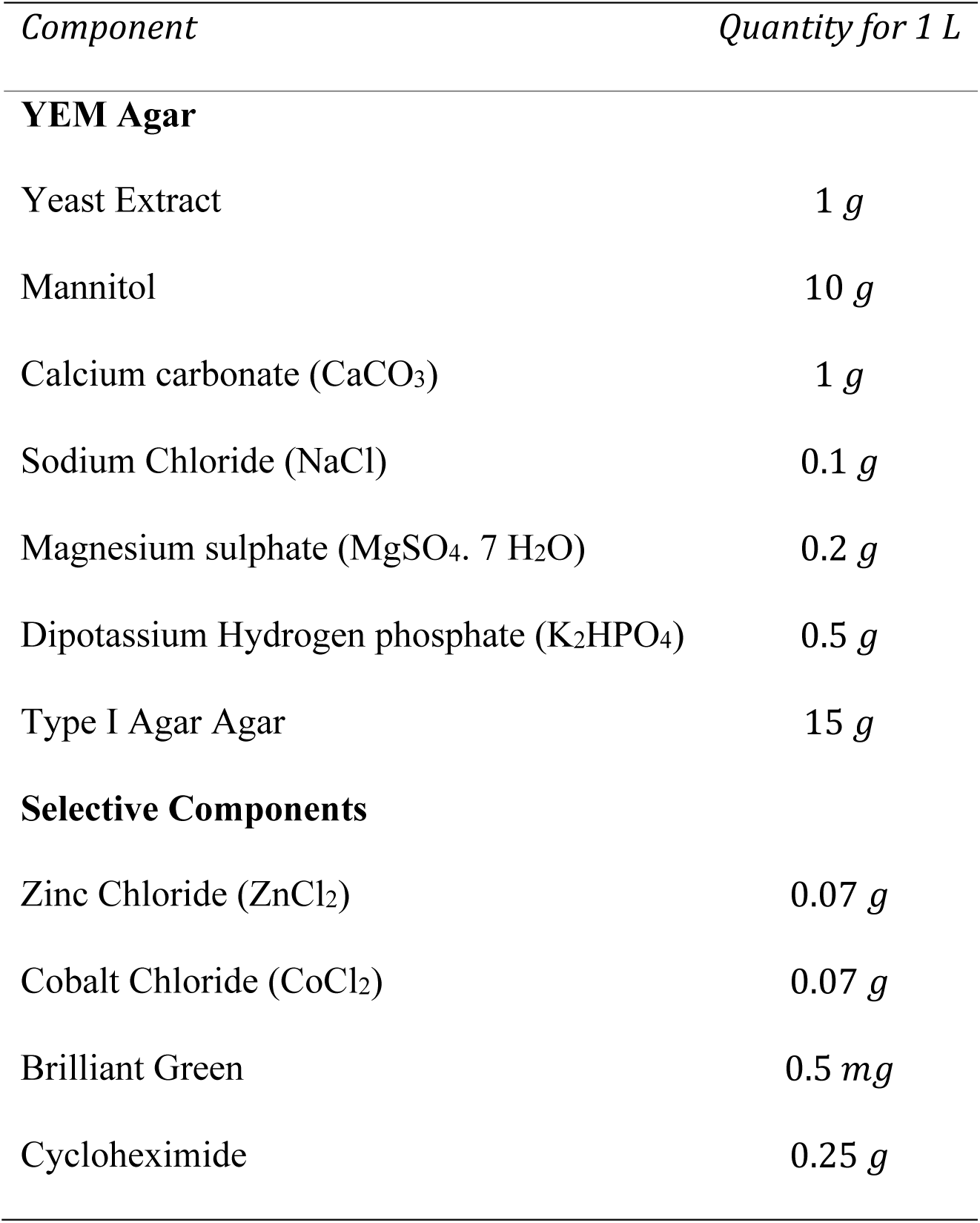
Composition of BRSM.

#### Selection of isolates

The colonies formed on the selective medium plates were examined, and the colonies that exhibit the characteristic features of *Bradyrhizobium sp.* were selected. The selected colonies were then inoculated in YEMA by quadrant streak technique to isolate single colonies. The single colonies thus isolated were grown in YEMB for about 6 to 7 days and stored.

### Confirmation of Isolated AINS-03 strains

The identity of the isolated *AINS-03* stains can be confirmed by performing various biochemical tests and staining techniques. Gram’s differential staining was performed to identify the Gram nature and to identify the morphological features. In addition, Bikrol, Saxena, and Singh (2010) conducted some Biochemical tests, including growth on Glucose-Peptone agar, Hofer’s alkaline medium, and Gelatin Liquefaction test Ketolactose production test and its action on milk.

#### Gram’s staining

Gram’s staining is a differential staining procedure that differentiates the bacteria into two broad groups, “Gram-Positive” and “Gram-Negative,” based on the cell membrane composition. Gram-Positive bacteria take up the primary stain while Gram-Negative bacteria take up the secondary stain.

The bacteria were stained with Crystal Violet, and then Mordent (Gram’s iodine) was added. Then the cells were decolorized with 95 % ethyl alcohol, and then the cells were stained using secondary stain Safranin and were observed under a microscope.

#### Growth on Glucose-Peptone agar

Glucose Peptone agar does not support the growth of Rhizobacteria (Pervin, Jannat, and Sanjee 2017). The Glucose Peptone agar was prepared (Table 2), sterilized, and poured into sterile plates, *AINS-03* isolated were inoculated by a simple streak technique, and the plates were incubated at 27 ℃ for eight days.

**Table 2.**
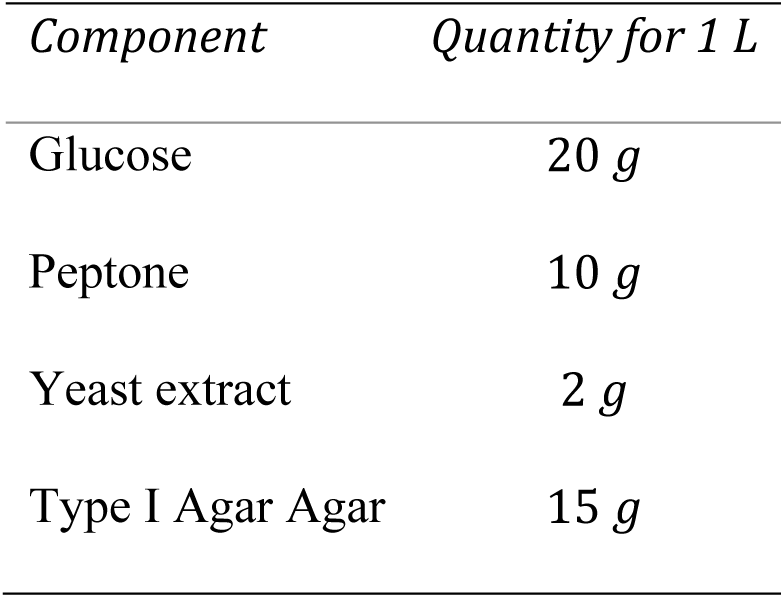
Composition of Glucose-Peptone agar.

#### Growth on Hofer’s Alkaline medium

The pH of Hofer’s alkaline medium is about pH 11, which does not support the growth of *Bradyrhizobium sp.* but other bacterial co-inhabitants of the rhizosphere, such as *Agrobacterium*. Therefore, to confirm that the isolated organism was *Bradyrhizobium sp.,* the organism was grown in Hofer’s alkaline medium.

Hofer’s alkaline medium (Hofer 1935) consists of all YEM Medium ingredients but Calcium Carbonate (𝐶𝑎𝐶𝑂_3_) and consists of Thymol blue as a pH indicator (Table 3). Isolated *AINS-03* strains were inoculated by a simple streaking technique and incubated at 27 ℃ for eight days.

**Table 3.** Composition of Hofer’s alkaline medium.

| *Component* | *Quantity for 1 L* |
| --- | --- |
| Yeast extract | 1 g |
| Mannitol | 10 g |
| Sodium Chloride (NaCl) | 0.1 g |
| Magnesium Sulphate (MgSO<sub>4</sub>. 7 H<sub>2</sub>O) | 0.2 g |
| Dipotassium Hydrogen phosphate (K<sub>2</sub>HPO<sub>4</sub>) | 0.5 g |
| Thymol Blue | 0.016 g |
| Type I Agar Agar | 15 g |

#### Gelatin Liquefaction test

*Bradyrhizobium sp*. does not liquefy gelatin. Gelatin medium (Thirst 1957) is prepared by mixing 0.1% peptone and 4% gelatin in distilled water and heating to 65 ℃, sterilized, cooled for 24 hours. Then the organism is inoculated by stabbing using inoculation needles and incubated at 26 ℃ for six days.

#### Keto lactose test

Some microorganisms produce keto lactose when inoculated in a medium containing lactose (Table 4). This property can be tested by inoculating the microorganism first in a glucose-containing medium and then in a lactose medium and then flooding the plate with Benedict’s reagent, which produces yellow colour when it reacts with keto lactose (JONES 1980).

**Table 4.** Medium for Keto lactose test.

| *Component* | *Quantity for 1 L* |
| --- | --- |
| **Medium 1** |  |
| Yeast extract | 10 *g* |
| Glucose | 20 *g* |
| Calcium carbonate (CaCO<sub>3</sub>) | 1 *g* |
| Type I Agar Agar | 15 *g* |
| **Medium 2** |  |
| Lactose | 10 *g* |
| Yeast extract | 1 *g* |
| Type I Agar Agar | 15 *g* |

This organism was grown in Medium 1 for about 24 – 48 hours and then transferred to plates with Medium 2 and grown for 24 – 48 hours. Then the plates were flooded with Benedict’s reagent and observed for colon changes.

#### Action on milk

The action of *Bradyrhizobium* strains on milk was studied by taking 15 mL of standardized cow milk and mixing it with 5 𝑚𝑙 of *AINS-03* strain with a cell concentration of about 108 𝑐𝑒𝑙𝑙𝑠/𝑚𝑙 CFU (Colony Forming Units). The mixture was swirled slowly and left undisturbed for about 15 minutes.

### Cultivation of Oryza sativa

#### Preparation of soil

Soil for the cultivation of *Oryza sativa* was taken from the lands of P. S. R. Engineering College, Sivakasi. The soil was sterilized using autoclave and dried. The soil was split into eight parts and filled in 8 pots, two of CP, RT, BT, and CF. 500 g of soil is taken and labelled as CTRL and analysed for initial NPK present in the ground.

#### Transplantation of seedlings

*Oryza sativa* seedlings of a variety of **white ponni** aged 40 days were obtained from a local farmer. The seedlings were grown on well-prepared land without applying any fertilizers of any kind. The seedlings were planted in a manner that each pot consisted of eight seedlings.

#### Fertilizer Treatments

Treatments were applied on the 10^th^ day and the 40^th^ day of transferring seedlings to pots. Pot named CP has not applied any treatment and is used as a control. The pot RT was applied with 1 𝑚𝑙/𝑙 of 10^8^𝑐𝑒𝑙𝑙𝑠/𝑚𝑙 CFU Liquid *Rhizobium* fertilizer obtained from Agricultural Service Centre, Thriuvengadam. The pot BT was applied with 1 𝑚𝑙/𝑙 of 10^8^𝑐𝑒𝑙𝑙𝑠/𝑚𝑙 CFU *AINS-03* strain grown in YEMB. The pot CF was applied with 50 𝑔/𝑙 chemical NPK complex fertilizer, recommended by the Agricultural Service Centre, Thriuvengadam.

#### Harvest

Grains produced were harvested after the 80^th^ day post-transplantation. Grains were then separated, dried, and packed in separate containers.

### Growth analysis of Oryza sativa

The growth of *Oryza sativa* is monitored by measuring four parameters viz., Plant height, Stem diameter, Leaf length, and leaf width. These parameters were measured after the seedlings were transferred to the pots with an interval of 5 days.

#### Plant Height

The height above the surface to the tip of the highest leaf was measured as Plant height. The height of the two plants in each pot was measured separately, and the mean was taken as the final plant height.

#### Plant diameter

The circumference of each plant was measured at a point 1 𝑖𝑛𝑐ℎ (25.4 𝑚𝑚) above the soil’s surface using a thread. Then, the diameter of the plant is measured by the formula.

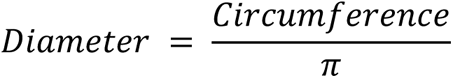

#### Leaf Length

Five random leaves were selected, and the length from the node to tip was measured. The mean of the length of leaves was calculated.

#### Leaf Width

The width of five random leaves at the broad portion of the leaf was measured. The mean of the width was calculated and recorded.

#### Grain Weight

The grains harvested were weighted using a digital balance. The mass of thousand grains was calculated by weighing one hundred random grains and multiplying the weight by 10.

### Soil Nutrition analysis

Soil nutrition enhanced by the *AINS-03* strain used was calculated by comparing the significant nutrients available in the soil before and after applying the four different treatments. Soils from all pots were collected after harvest. Then they were dried, powdered, and packed in packets labelled appropriately and stored. Soil before treatment was collected initially and labelled as CTRL.

Available significant nutrients in the soil such as Nitrogen, Potassium, and Phosphate were analysed by availing the Soil test service of “Krishi Vigyan Kendra,” Aruppukottai, Virudhunagar (Dt.) (KVK Virudhunagar).

#### Nutritional Value of Oryza sativa

Food is composed of three major nutrients such as carbohydrates, fat, and proteins. Therefore, by estimating the amount of total carbohydrates, fats, and proteins, we can find the energy provided by rice.

Rice samples were boiled, and the husk was separated and dried. Then, the endosperm was powdered, and the solution containing 1 𝑚𝑔/𝑚𝑙 of rice sample was prepared and stored as stock solution.

#### Estimation of Total Carbohydrates

Total carbohydrates present in the rice are estimated by using the Anthrone method. First, carbohydrates are dehydrated by concentrated sulphuric acid to formal Furfural. Then, Furfural condenses with Anthrone to form a blue-coloured complex which can be measured calorimetrically (Yemm and Willis 1954).

To 1 𝑚𝑙 10% stock solution, 4 𝑚𝑙 of Anthrone reagent was added and mixed well and incubated in boiling water for 10 minutes then cooled and optical density was measured at 630 𝑛𝑚 using UV-Visible spectrophotometer. Glucose solution was used as a standard for the estimation of total carbohydrates present in the sample.

#### Estimation of Total Proteins

Total proteins present were estimated by the method described by Lowry et al. (1951). Under alkaline conditions, proteins react with copper sulphate to form a copper-protein complex. When reacted with the Folin Ciocalteau Phenol reagent, this complex reduces phosphor molybdenum and phosphor tungstate to heteropoly molybdenum and tungstic acid. Aromatic amino acids such as tyrosine and tryptophan react with those compounds and produce a dark blue complex that obeys Beer-Lamberts law at 520 𝑛𝑚.

Sample solutions were prepared by taking 10% solution. To 1 𝑚𝑙 of sample solution, the alkaline copper solution was added and incubated at room temperature for 10 minutes. Then 0.5 𝑚𝑙 of Folin Ciocalteau Phenol reagent was added and incubated at room temperature for 10 minutes. Optical density was measured at 520 𝑛𝑚. Bovine Serum Albumin (BSA) was used as standard, and the concentrations of proteins present in samples were found.

#### Estimation of Total Lipids

Total lipids present in the grains were estimated by a method described by Rawle, Anderson, and Knight (1972). Lipids react with vanillin in a medium containing concentrated sulphuric acid and phosphoric acid. Concentrated sulphuric acid reacts with unsaturated lipids to form a carbonium ion. Phosphoric acid reacts with vanillin to produce phosphate ester, which increases the reactivity of the carbonyl group. The carbonium ions react with the carbonyl group and form a coloured compound whose optical density can be read at 525 𝑛𝑚.

Sample solutions were prepared by taking 10% stock solution. To 0.1 𝑚𝑙 of sample solution, 5 𝑚𝑙 of concentrated sulphuric acid were added and incubated in a boiling water bath for 10 minutes. Then 0.4 𝑚𝑙 of aliquot was transferred to a clean, dry tube. 6 𝑚𝑙 of Phosphor-Vanillin reagent was added to the aliquot and incubated in the dark for 45 minutes. Optical density was measured at 525 𝑛𝑚. 1 𝑔/𝑙 olive oil in absolute ethanol was used as standard, and 0.4 𝑚𝑙 concentrated sulphuric acid as blank.

## Result and discussion

### Isolation of AINS-03 strains

*AINS-03* strains that were grown in the selective medium plates were observed. Colonies formed in the plate spread with 10^-5^ and 10^-6^ dilutions were too many to count. Eight single colonies with the same morphology were found in the plate spread with 10-7 dilution. In the plate with 10^-8^ dilution, no colonies were found.

Eight colonies of the plate with 10^-7^ dilution were plated separately in the BRSM using the quadrant streak technique to isolate single colonies. The colonies from the 3^rd^ plate (AINS-03) shared the same morphology of *Bradyrhizobium japonicum* strains described in Bergey’s Manual of Determinative Bacteriology (Bergey and Breed 1957). *Bradyrhizobium japonicum* is the type strain for *Bradyrhizobium* genus.

### Confirmation of Isolated AINS-03 strains

The characteristics of AINS-03 strains isolated were confirmed by comparing it with the reference strain MCC3320 obtained from National Centre for Microbial Resource (NCMR), Pune, India. The comparison of characteristics of isolated and reference strains were presented in table 5.

**Table 5.** Comparison of characteristics of isolated and References Strain MCC3320.

| *Test* | *Reference Strain*<br>*(MCC3320)* | *Isolated Strain*<br>*(AINS-03)* |
| --- | --- | --- |
| Gram Staining | -ve | -ve |
| Growth on Glucose-Peptide agar | -ve | -ve |
| Growth on Hofer's alkaline medium | -ve | -ve |
| Gelatin Liquefaction | -ve | -ve |
| Ketolactose production | -ve | -ve |
| Action on Milk | Curdling | Curdling |

No growth was found on the Glucose-Peptone agar and Hofer’s alkaline medium even after 10 days after inoculation, which is the expected period of growth to achieve significant colony size. The same was observed on the Gelatin liquefaction test also. No liquefaction was observed after 10 days of inoculation.

Ketolactose production test was performed by inoculating the AINS-03 strains in “Medium 1” in a test tube slant and incubated at 27 ℃ for about 48 hours. Only a small growth was observed and then streaked on “Medium 2” and incubated for 48 hours. Plates were then flooded with Benedict’s reagent. No colour change was observed after 1 hour. When AINS-03 strains were added to milk, Curdling was observed after 5 minutes. The amount of curdles formed increased with time. Reference strain *Bradyrhizobium sp.* (MCC3320) strain produced a high amount of curdling while the isolated strain *Bradyrhizobium sp*. (AINS-03) had a slightly lesser curdling than strain MCC3320. All the results obtained were on par with the previous results obtained by Bikrol, Saxena, and Singh (2010) and with Bergey’s Manual of Determinative Bacteriology (Bergey and Breed 1957).

### Cultivation of Oryza sativa

The soil was prepared in a suitable condition with appropriate amounts of soil and water two days before the transplantation of seedlings. Seedlings obtained were planted morning. On the day of the plantation, the height of each plant was measured. Plants planted in the pot BT started to flower on the 53^rd^ day after transplantation, followed by the plants on pot CF on the 57^th^ day. CP and RT blossomed on the 63^rd^ day after transplantation. After the 80^th^ day, the grains were harvested.

### Growth analysis of Oryza sativa

#### Plant Height

The plant height of Oryza sativa was measured once every five days. The day of transplantation was marked as the 0^th^ day, and the height of the plant above the surface was about 17.1 to 17.3 cm. Plants in all pots showed a similar growth till the 10^th^ day. Variations in height were observed after the applications of fertilizer treatments. Plants applied chemical fertilizers (CF) showed a difference from day 15 onwards. It showed maximum growth than the other three till the end. Plants applied with AINS-03 strain (BT) followed those applied with chemical fertilizers with second maximum growth. Plants applied with Rhizobium strains (RT), and control (CP) showed similar growth lesser than CF and BT (Figure 1).

**Figure 1.**
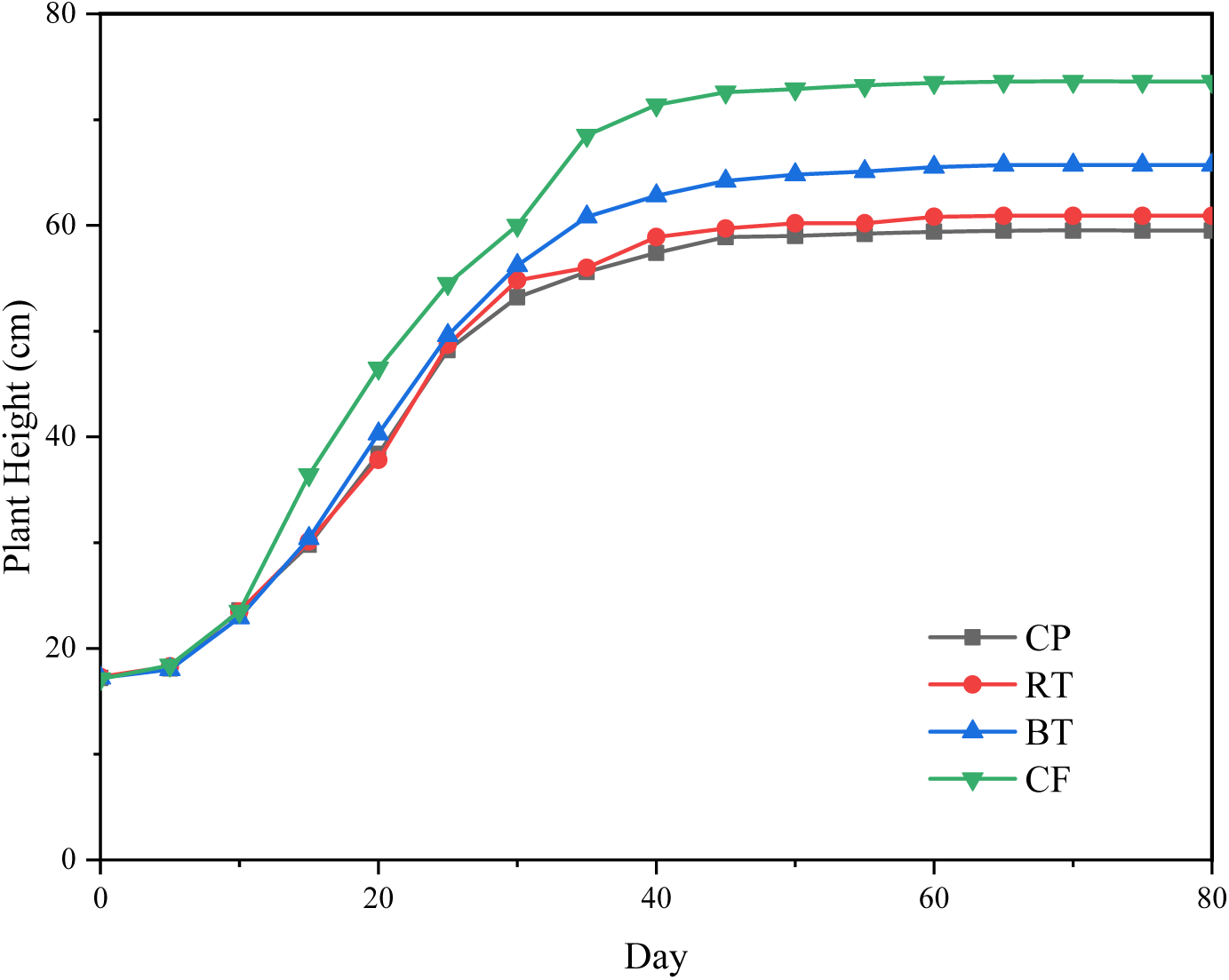
Graph showing plant height in different treatments

Plants in pots CP and RT showed a close growth, with RT showing slightly higher growth than CP. BT seems to have grown like CP and CF until the 25th day, but the height increases gradually and makes a significant difference from CP and RT. The growth of all plants stopped after the first flower appeared on each plant. Grains started to show after five days of flowering. It took almost ten days for a plant to produce flowers and grains on every stalk. CF reached a maximum height of 73.6 cm, BT reached 65.7 cm, RT reached 60.9 cm, and CP reached 59.5 cm.

#### Plant Diameter

The diameter of the plant was measured by using a thread. The thread was wrapped around the stem above one inch from the ground to measure the circumference of the stem. From the circumference, radius and diameter are calculated. The initial diameter of the plant was about 4.5 to 5.0 mm. Like Plant height, all plants have a similar diameter till day 15. After day 15, the diameters started to accelerate. CF showed the highest diameter, which BT follows. The plant height of CP and RT is almost the same, but in diameter, RT shows a significant increase. The increase in diameter has stopped only after the 70^th^ day (Figure 2).

**Figure 2.**
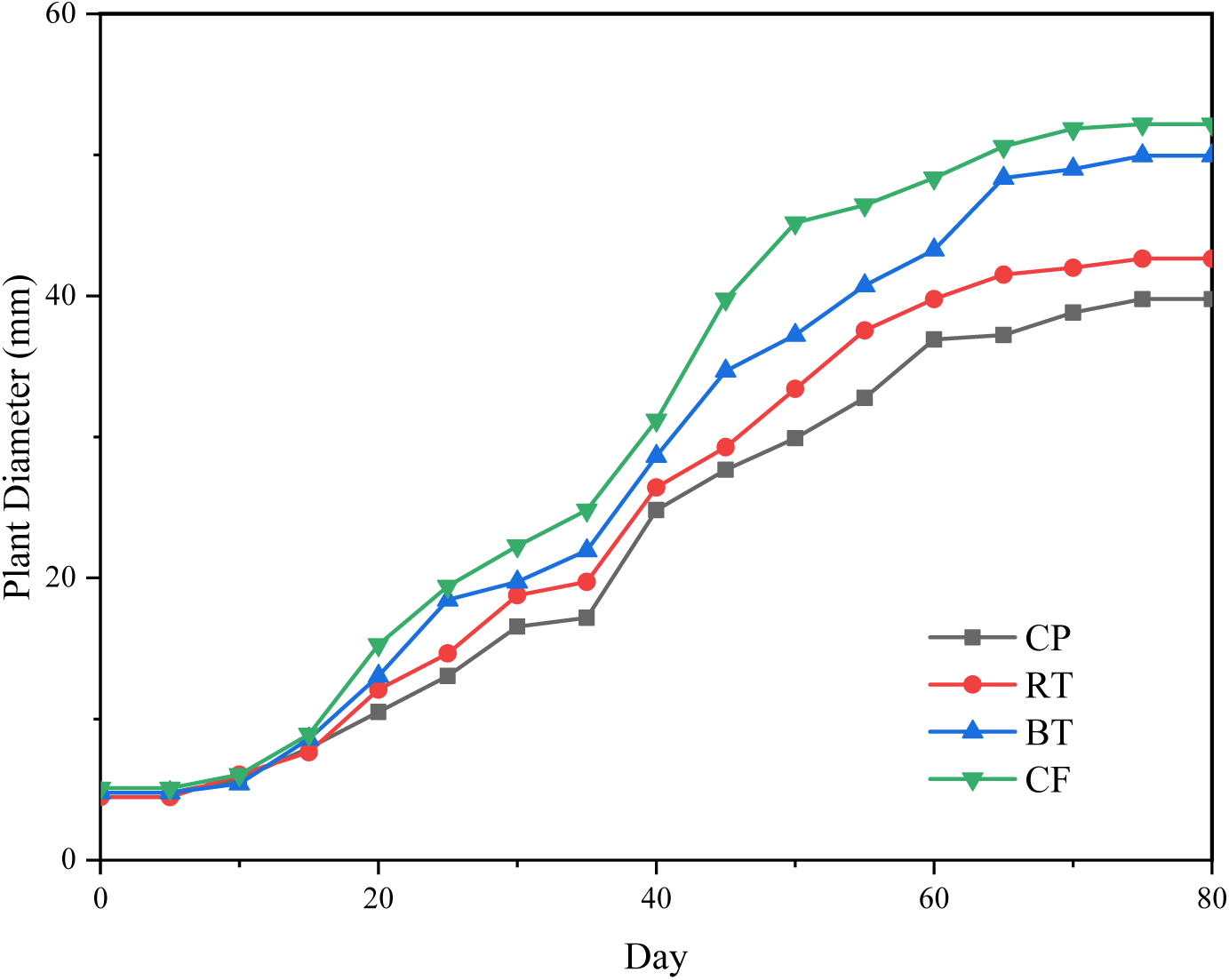
Graph showing plant diameters in different treatments

Before reaching a constant, the diameter of BT increases at a steady rate and gets close to CF on the 65th day, which we believe due to the grains were emerging and thickening during that period. CP had the final diameter of 39.77 mm, RT had 42.64 mm, BT had 49.95 mm, and CF had 52.18 mm.

#### Leaf Length

Leaf Length was measured from node to tip of a leaf. The initial average length of leaves was about 10.20 cm to 11.48 cm. The size of leaves showed a close similarity till the 15^th^ day. After the 15^th^ day, the leaf length of CF increased significantly from other treatments. Length of BT, RT, and CP was similar till the 25^th^ day. The length of leaves almost stopped growing after the 60^th^ day. Unlike in the height and diameter of plants, the leaves always showed a close difference in length and not a significant variation. Leaves of CF showed a sudden acceleration in length around day 35 and followed a constant increase (Figure 3).

**Figure 3.**
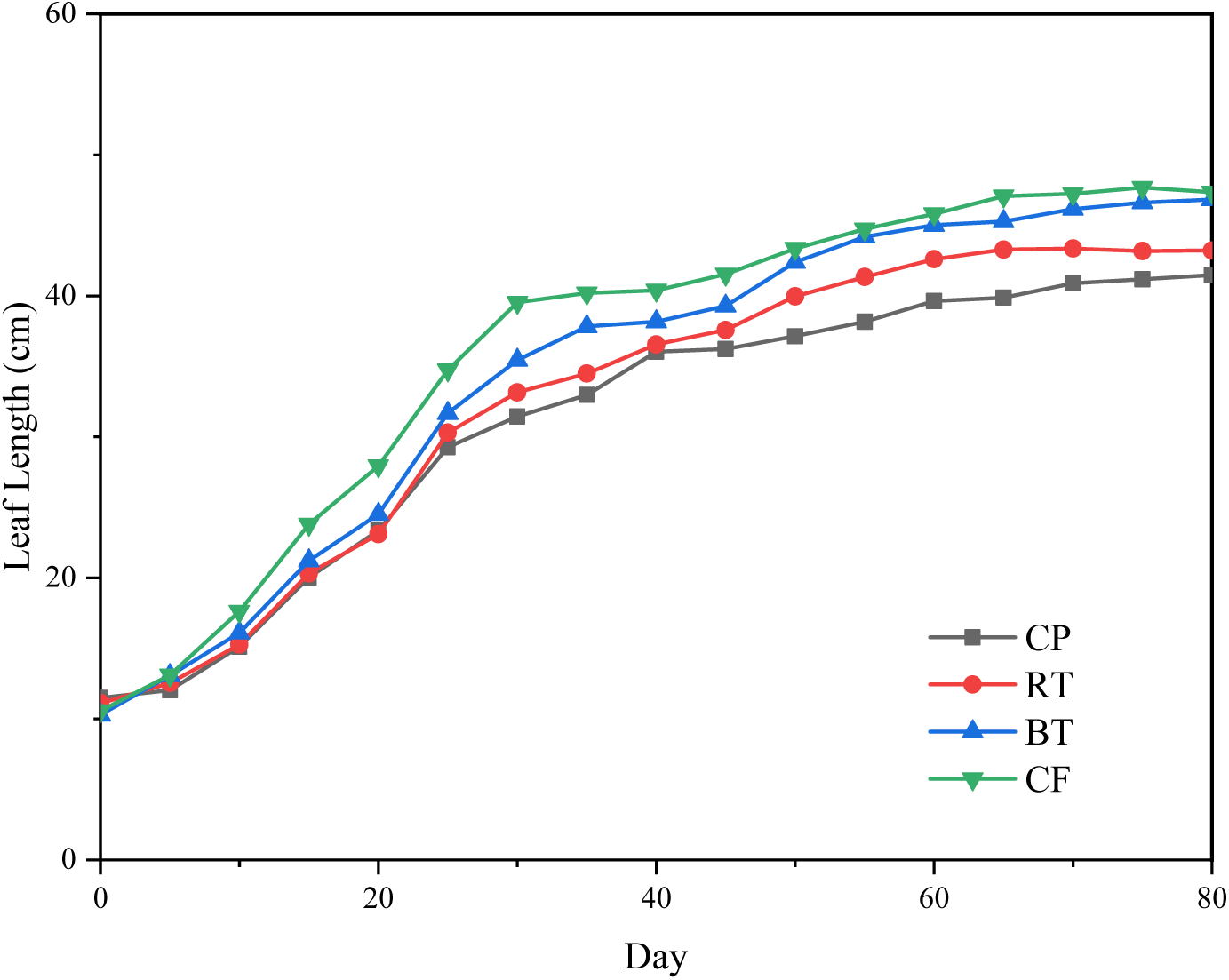
Graph showing Leaf length in different treatments

While the length of leaves of BT followed a constant rate of increase, and in the end, they almost came close to those of CF. Leaves of CP and RT showed a close rate till day 50. After that, there was a significant increase in the length of leaves of RT and remained constant after day 65. Thus, leaves CP and RT ended up in almost close lengths. The highest average leaf length observed in CP was 41.8 cm, RT was 43.22 cm, BT was 46.84 cm, and CF was 48.36 cm.

#### Leaf Width

Leaf width was measured at the broad portions of the leaf. The average Initial leaf width was in the range of 0.28 mm and 0.32mm. Leaf width in the four pots was almost the same till day 10. After the applications of treatments, the leaf width showed a significant difference. As in other parameters, CF showed a maximum leaf width on all days, followed by BT. Leaves of CF showed an increase in width from other treatments on day 15, while the other treatments started to show a difference only after the 25^th^ day. The width of CF increased at a greater rate. The width of BT and RT showed a difference from CP on the 30^th^ day onwards (Figure 4).

**Figure 4.**
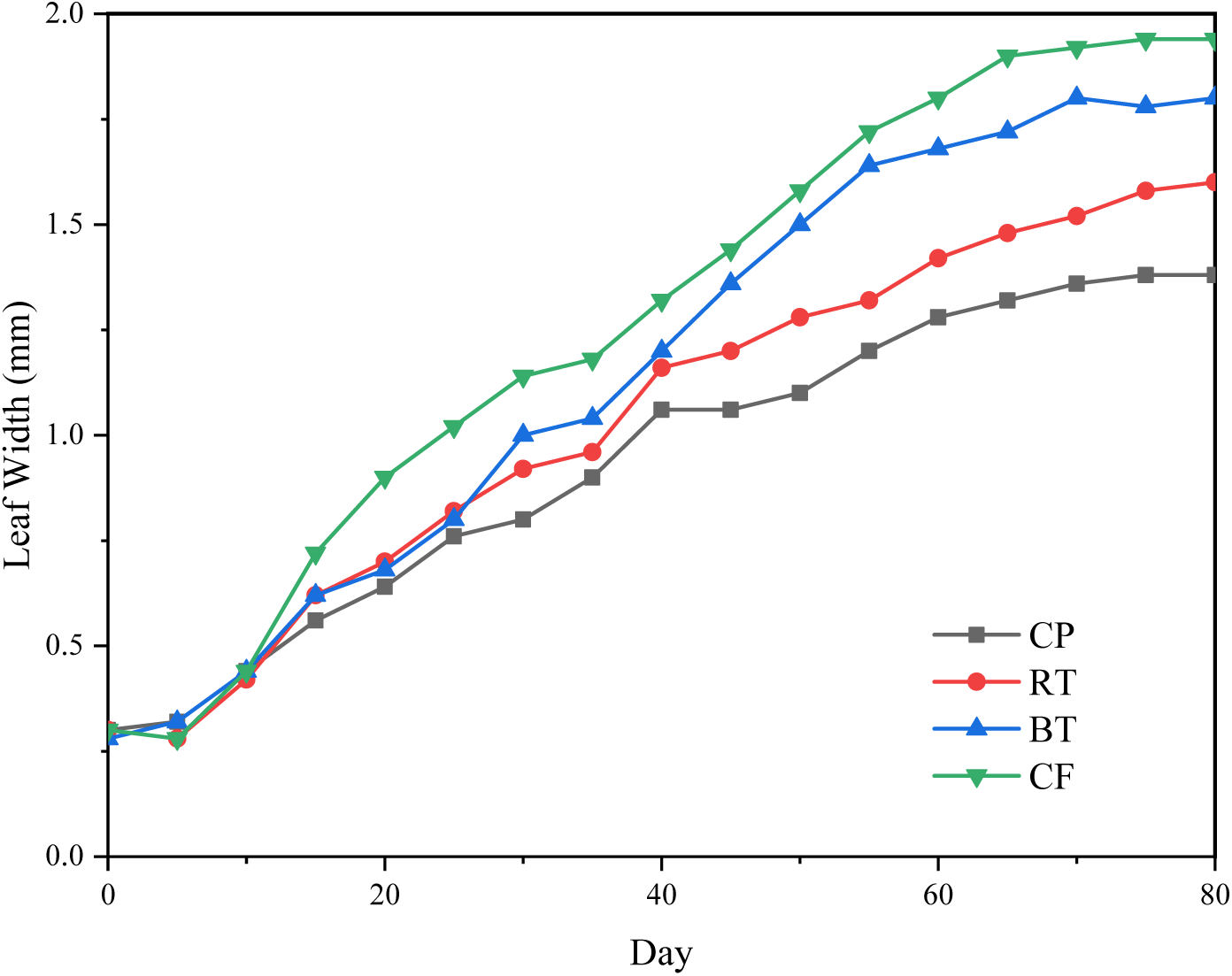
Graph showing the leaf width in different treatments

Leaf width of CF increased at a constant rate till day 65 and became constant afterward. After the 30^th^ day, the width of BT leaves started to show a steady rate of increase till Day 50. Then, the rate decreased and became constant after day 70. Leaf width of CP and RT were similar till day 25, and then the rate increased for RT leaves till day 40. After day 40, the leaf width of RT increased at a constant rate till the end. Also, the width of CP leaves showed a constant increase till day 40 but at a lesser rate than RT. Then, the rate decreased much for the next ten days and started to increase afterward. The increase in leaf width of CP stopped after day 65 and remained constant. The final leaf width observed in plant CP was 13 cm, RT was 1.6 cm, BT was 1.8 cm, and CF was 1.94 cm.

#### Grain Weight

Grains collected were dried, and the weight of 100 grains was measured using physical balanced and the total yield was also measured (Table 6). The average weight of 1000 grains of White Ponni variety were 16 g, as indicated by the Tamil Nadu Agricultural University (TNAU) website, Agritech portal (Bergey and Breed 1957). The weight of grains from CP was about 14.1 g which is 1.9 g less than the average weight recommended. RT weighted 14.4 g for 1000 grains which are 1.4 g less than the average weight. BT and CF had weights higher than the average weight. BT had 18.2 g, 2.2 g higher than the average weight, and CF had 17.5 g, which is 1.75 g higher than the average weight for 1000 grams. The growth of CF plants is higher than that of BT, which is also reflected in the total yield. CF had the maximum yield, followed by BT. CP and RT had almost a similar yield.

**Table 6.** Grain weight of Different treatments.

| *Treatment* | *Weight of 100*<br>*grains (g)* | *Weight of 1000*<br>*grains (g)* | *Total Yield (g)* |
| --- | --- | --- | --- |
| CP | 1.41 | 14.1 | 34.3 |
| RT | 1.44 | 14.4 | 34.5 |
| BT | 1.82 | 18.2 | 37.2 |
| CF | 1.75 | 17.5 | 38.4 |

Physical observation of grain size revealed the grain size of BT is slightly higher than CF. Although the size of grains also impacted the weight of grains, BT produced grains with the highest weight, followed by CF. Grains from RT and CP had almost similar weight, but RT had a slightly higher grain weight than CP.

### Soil Nutrition Analysis

Soil Nutrition analysis was made by availing the service of KVK, Virudhunagar. Total available nitrogen after cultivation was compared with initially available nitrogen which was 160 kg ha^-1^. Soil that is applied with chemical fertilizers (CF) shown a maximum nitrogen content of about 196 kg ha Soil applied with AINS-03 strains (BT) contained 192 kg ha^-1^ of nitrogen which is very close to that of soil treated with chemical fertilizer CF. Rhizobium treated soil RT had a minimal nitrogen fixed 170 kg ha, which is only 10 kg ha^-1^ higher than the initial available amount. Soil without any treatment CP consisted of 166 kg ha^-1^ of total available nitrogen (Table 7).

**Table 7.** Total available nitrogen in soil.

| *Treatment* | *Total Available Nitrogen (kg ha<sup>-1</sup>)* |
| --- | --- |
| Initial (CR-I) | 160 |
| CP | 166 |
| RT | 170 |
| BT | 192 |
| CF | 196 |

From the above results, the nitrogen fixed by AINS-03 strains in the soil is very close to that of chemical fertilizers. Also, the nitrogen fixed by Rhizobium strains is very low. This is due to the symbiotic nature of the Rhizobium. As the crop is cereal, there is no way of integrating with the root. However, some rhizobium strains may colonize the root for a while and fix some nitrogen.

### Nutritional Value of Oryza sativa

Total available carbohydrates, proteins, and fat in the grains obtained from four different treatments were estimated (Table 8). The energy provided by each grain can be calculated by knowing the energy supplied by each essential macronutrient. Each gram of Carbohydrate can provide four calories of energy. Each gram of Protein can provide four calories of energy. Each gram of fat can provide nine calories of energy.

**Table 8.** Nutritive value of Oryza sativa and energy obtained (Per 100 g of rice)

| *Treatment* | *Carbohydrate*<br>*(g)* | *Protein (g)* | *Fat (g)* | *Energy (cal)* |
| --- | --- | --- | --- | --- |
| CP | 53.7 | 6.1 | 0.23 | 241.63 |
| RT | 54.5 | 6.7 | 0.26 | 247.14 |
| BT | 66.8 | 8.1 | 0.27 | 302.03 |
| CF | 63.1 | 7.8 | 0.28 | 286.12 |

Grains from plants treated with AINS-03 strain (BT) had the highest nutritional value and energy compared to other treatments. Plants applied with chemical fertilizers (CF) produced grains with a nutritional value close to the BT. Grains from plants applied with Rhizobium strains had lower nutritive value than CF but higher than the grains from plants without any fertilizer treatment (CP).

Protein content in the plants’ BT and CF is very high than in the other treatments. On the other hand, the fat content available in plants treated with Rhizobium seems to be increased than the control, as the other nutrients were in a similar composition.

## Conclusion

*Bradyrhizobium* species isolated from *Aeschynomene indica*’s rhizosphere have the free-living ability and plant growth-promoting ability. Also, these bacteria are known to synthesize plant growth hormones such as Gibberellins and IAAs, which may be the cause here in plant growth promotion.

In the present study, AINS-03 strains were tested for nitrogen-fixing and plant growth promotion on cereals. Soil treated with AINS-03 strains showed the highest nitrogen fixation of 192 kg ha^-1,^ which is slightly low than the nitrogen in chemical fertilizer treated soil (196 kg ha^-1^).

Plant growth promotion by AINS-03 is very similar to the growth promoted by chemical fertilizer. In growth, parameters observed such as Plant height, diameter, leaf length, and leaf width, plants treated with AINS-03 are less than those treated with chemical fertilizer. Despite low growth than plants applied chemical fertilizer, AINS-03 treated plants produced a better yield and grains with high nutritional value than others.

Rhizobium can be used as a PGPR only for legumes because of its symbiotic nature and host specificity. As AINS-03 strains are capable of free-living in the soil, we propose that the AINS-03 strains be used as nitrogen fixers and Plant growth-promoting plant rhizobacteria for both Pulses and Cereals. Further research can be made to optimize the method of applying fertilizer, time, and frequency of fertilizer treatment with AINS-03 sp. as Bio-Fertilizer.

## Notes

### Competing Interest Statement

The authors have declared no competing interest.

